# Phylogeographic structure of the pygmy shrew: revisiting the roles of southern and northern refugia in Europe

**DOI:** 10.1101/819912

**Authors:** Rodrigo Vega, Allan D. McDevitt, Joanna Stojak, Alina Mishta, Jan M. Wójcik, Boris Kryštufek, Jeremy B. Searle

## Abstract

Southern and northern glacial refugia are considered paradigms that explain the complex phylogeographic patterns and processes of European biota. Although the Eurasian pygmy shrew *Sorex minutus* Linnaeus, 1766 (Eulipotyphla, Soricidae) has been used a model species to study geographic isolation and genetic diversification in Mediterranean peninsulas in the Last Glacial Maximum (LGM), and post-glacial population expansion from cryptic northern glacial refugia in Western and Central Europe, there has been incomplete knowledge about the phylogeographic structure, genetic differentiation and demographic history within these regions. Here, we provide a revisited statistical phylogeographic study of *S. minutus* with greater sampling coverage in terms of numbers of individuals and geographic range, making it the most comprehensive investigation of this species to date. The results showed support for genetically distinct and diverse phylogeographic groups consistent with southern and northern glacial refugia, as expected from previous studies, but also identified geographical barriers concordant with glaciated mountain ranges during the LGM, early diversification events dated between the Upper Pleistocene and Lower Holocene for the main phylogeographic groups, and recent (post-LGM) patterns of demographic expansions. The results have implications for the conservation of intraspecific diversity and the preservation of the evolutionary potential of *S. minutus*.

## INTRODUCTION

During the Quaternary glaciations, species in Europe were restricted to glacial refugia at glacial maxima (Bilton *et al*., 1998; Taberlet *et al*., 1998; Hewitt, 2000; Stewart & Lister, 2001; Pazonyi, 2004; Sommer & Nadachowski, 2006). As glaciers retreated, a broad range of recolonisation patterns emerged, as evidenced by palaeontological, biogeographic and phylogeographic studies on various taxa, resulting in the complex contemporary patterns of endemism, species richness and biodiversity hotspots observed across Europe. While population contraction and lineage diversification within southern glacial refugia in the Mediterranean peninsulas during the Last Glacial Maximum [LGM; 19-26.5 thousand years ago (KYA) (Clark *et al*., 2009)], and subsequent northward postglacial recolonisation of Europe have been accepted and recognised since the 1990s (Bilton *et al*., 1998; Taberlet *et al*., 1998; Hewitt 2000), the concept of cryptic northern glacial refugia also became a paradigm to explain the complex phylogeographic patterns and processes of European biota (Stewart & Lister, 2001; Pazonyi 2004; Sommer & Nadachowski, 2006). Fossil records and phylogenetic analyses revealed that many species of flora and fauna could have survived during the LGM in the Carpathian Basin (Stewart & Lister, 2001; Pazonyi, 2004; Sommer & Nadachowski, 2006; Stojak *et al*., 2015), in Dordogne region (Steward *et al*., 2010) and in the Ardennes (Stewart & Lister, 2001), and glacial refugia could also be located in Crimea (Marková, 2011) or the Russian Plain (Banaszek *et al*., 2012). Nowadays, locations of southern and northern glacial refugia during the LGM are hotspots of genetic diversity (Petit *et al*., 2003; Stojak *et al*., 2016).

The Eurasian pygmy shrew *Sorex minutus* Linnaeus, 1766 (Eulipotyphla, Soricidae) (Hutterer, 1990) has been used as a phylogeographic model species for studying the persistence of populations in northern European refugia during the LGM (Bilton *et al*., 1998: McDevitt *et al*., 2010; Vega *et al*., 2010a, b). It is one of the few mammalian species that is widely distributed in the three Mediterranean peninsulas, and in Central and Northern Europe (Fig. 1A); therefore, *S. minutus* is an excellent model for understanding the effects of the glaciations in Europe and the colonisation history during the Pleistocene and postglacial times. Although several studies have found evidence supporting the hypotheses of southern geographic isolation and genetic diversification, and population expansion from cryptic northern glacial refugia (Bilton *et al*., 1998; McDevitt *et al*., 2010; Vega *et al*., 2010a, b), little is known about the phylogeographic structure, genetic differentiation and demographic history of this small mammal within these regions due to the limited number of samples from Mediterranean peninsulas. An expanded phylogeographic study of the pygmy shrew is therefore important for the understanding and further development of biogeographic models of glacial refugia and postglacial recolonization, for depicting areas with high intraspecific genetic diversity, for establishing conservation measures of rear-edge populations, and for the preservation of the evolutionary potential of species, particularly in the face of climate and anthropogenic change (Deffontaine *et al*., 2005; Provan & Bennett, 2008; Stojak *et al*., 2019; Stojak & Tarnowska, 2019).

**Figure 1.**
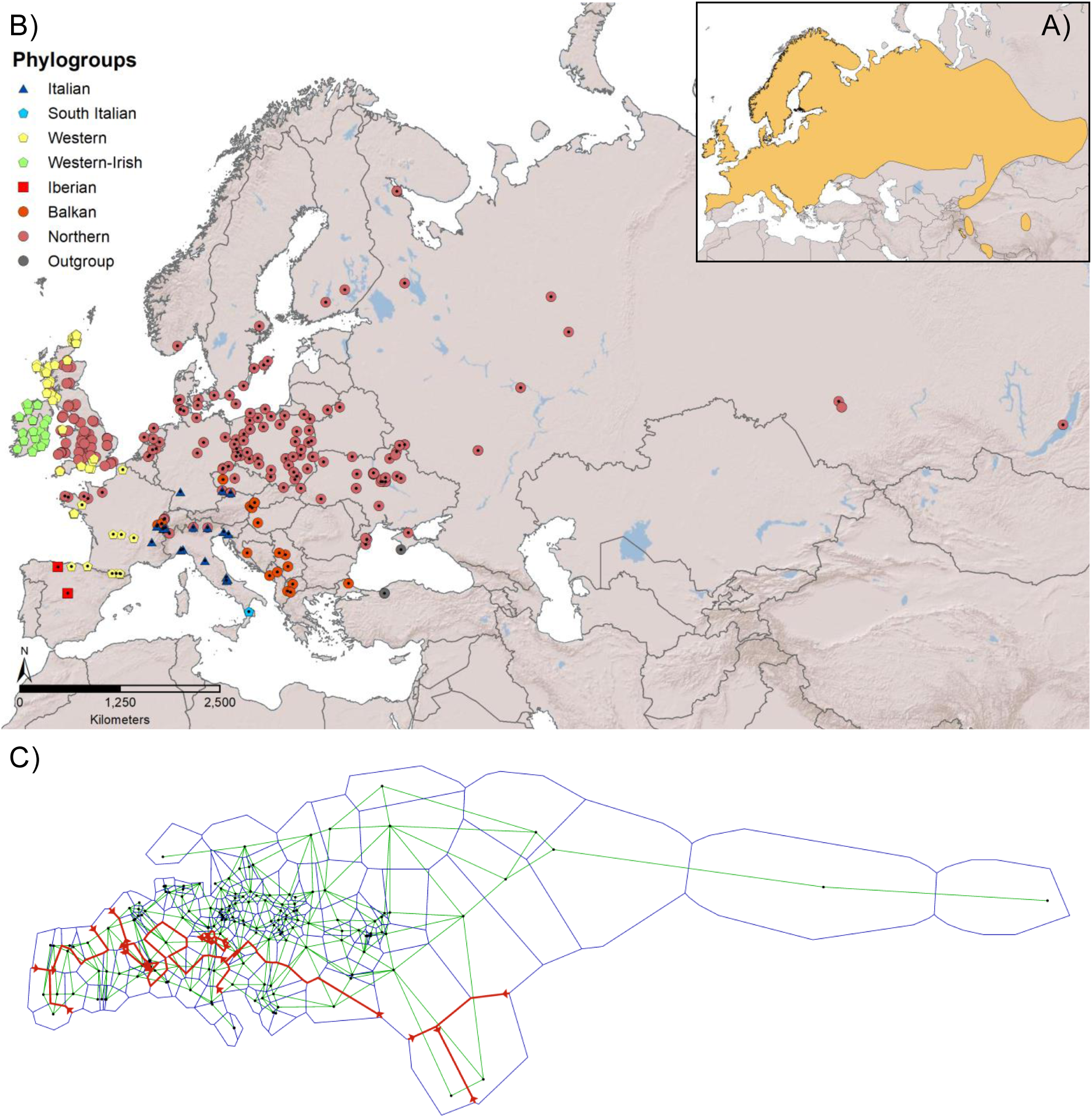
A) Map of Eurasia showing the geographical distribution of the Eurasian pygmy shrew *Sorex minutus* (Hutterer *et al*., 2016). B) Sample localities of *S. minutus* used for this study and divided into cytochrome (cyt) b phylogroups (symbols with a dot represent samples used for inferring geographic barriers). C) Geographic barriers (red lines) for *S. minutus*; the barriers (up to a maximum of 10) were inferred using Monmonier’s maximum difference algorithm which finds edges (boundaries) on the Voronoi tessellation (blue polygons) associated with the highest rate of change in genetic distances among a subset of continental samples (dots) interconnected with a Delaunay triangulation (green lines).

In this study, we explored the evolutionary history and phylogeographic structure of *Sorex minutus* using a statistical phylogeography approach (Knowles & Maddison, 2002; Knowles, 2009). Here, we emphasise the genetic diversity and structure within and among refugia, the inference of geographical barriers and the demographic history of *S. minutus*, which are aspects that have not been studied in detail previously. Specifically, we asked the following questions: 1) What are the geographical distribution and genetic diversity patterns of the genealogical lineages of *S. minutus*? 2) Is there an isolation-by-distance pattern across the geographic range of *S. minutus* or do the lineages show significant population genetic structure? 3) What is the historical demography of *S. minutus* in Europe? Our results showed support for distinct and genetically diverse lineages, geographical barriers concordant with glaciated mountain ranges during the LGM, and recent (post-LGM) population expansions with contemporary contact areas. The results presented here have implications for the long-term conservation of intraspecific diversity and the preservation of the evolutionary potential of *S. minutus* in the face of modern climate change.

## MATERIALS AND METHODS

### Study species

*Sorex minutus* is common over most of its distribution but is rarely dominant and it occurs in a wide range of terrestrial habitats with adequate ground cover and in relatively damp areas, including swamps, grasslands, heaths, sand dunes, woodland edge, rocky areas, shrubland and montane forests (Hutterer, 1990, 2016; Churchfield, 1990; Churchfield & Searle, 2008). It is found from southern and western Europe to much of central and northern Europe, Ireland and the British Isles, and Siberia to Lake Baikal in the east (Hutterer, 1990, 2016). It is found from sea level up to 2260 m (in the Alps), but its distribution becomes patchy and limited to higher altitudes in southern Europe where it occurs with some degree of geographical isolation and differentiation, while in central and northern parts of Europe and in Siberia it is more abundant and populations are more connected and widespread (Hutterer, 1990, 2016).

### Samples and molecular methods

A total of 671 cytochrome b (cyt b) DNA sequences of *S. minutus* from Europe and Siberia were used for this study (Fig. 1B; see Supplementary information Table S1). DNA sequences were obtained from samples collected from the wild following ethical guidelines (Sikes, Gannon & the Animal Care and Use Committee of the American Society of Mammalogists, 2011), or from museums, and from published GenBank data (including AB175132: Ohdachi *et al*., 2006; AJ535393 – AJ535457: Mascheretti *et al*., 2003; GQ272492 – GQ272518: Vega *et al*., 2010a; GQ494305 – GQ494350: Vega *et al*., 2010b; and JF510376 – JF510321: McDevitt *et al*., 2011). In addition, four cyt b sequences of *S. volnuchini*, which was used as an outgroup (Fumagalli *et al*., 1999), were incorporated into the analysis (including AJ535458: Mascheretti *et al*., 2003).

Genomic DNA from wild and museum samples was extracted using a commercial kit (Qiagen). Partial (1110 bp) cyt b sequences were obtained by PCR using two primer pairs that amplified approximately 700 bp of overlapping fragments, or using five primer pairs (for museum samples with highly degraded DNA) that amplified approximately 250 bp of overlapping fragments (Vega *et al*., 2010a). PCR amplification was performed in a 50 μl final volume: 1X Buffer, 1 μM each primer, 1 μM dNTP’s, 3 mM MgCl_2_ and 0.5 U Platinum Taq Polymerase (Invitrogen), with cycling conditions: 94°C for 4 min, 40 cycles at 94°C for 30 s, 55°C for 30 s and 72°C for 45 s, and a final elongation step at 72°C for 7 min. Purification of PCR products was done with a commercial kit (Qiagen) and sequenced (Macrogen and Cornell University Core Laboratories Center).

### Phylogenetic analysis

Sequences were edited in BioEdit version 7.0.9.0 (Hall, 1999), aligned in ClustalX version 2.0 (Larkin *et al*., 2007). A haplotype data file was obtained using DnaSP version 5.10.1 (Librado & Rozas, 2009). Newly obtained haplotypes were deposited in GenBank (Accession Numbers: XXXXX – XXXXX).

The model of evolution that best fitted the molecular data (haplotypes) was searched using jModelTest version 2.1.10 (Darriba *et al*., 2012) using the Bayesian Information Criterion value. The substitution model supported was the GTR with specified substitution types (A–C=0.4250, A–G=23.5124, A–T=1.6091, C–G=1.8671, C–T=17.2314, G–T=1.0000), proportion of invariable sites (0.6044), gamma shape parameter (0.2816) and nucleotide frequencies (A=0.2777, C=0.3076, G=0.1416, T=0.2731).

The phylogenetic relationships among cyt b haplotypes of *S. minutus* were inferred by Bayesian analysis and by generating a parsimony phylogenetic network. The Bayesian analysis was done using MrBayes version 3.2.7 (Ronquist *et al*., 2012) with two independent runs (10 million generations and 5 chains each), a sampling frequency every 1000 generations and temperature of 0.1 for the heated chain, and checking for convergence in Tracer version 1.7.1 (Rambaut *et al*., 2018). Trees were summarized after a burn-in value of 2500 to obtain the posterior probabilities of each phylogenetic branch. The main phylogenetic groups (phylogroups) were identified based on monophyly of the haplotypes, and were named based on the geographical origin of the samples. The phylogenetic network was done using PopART version 1.7 (http://popart.otago.ac.nz) implementing a median-joining algorithm.

Sequence polymorphism indices and diversity values, including the number of haplotypes (H), polymorphic (segregating) sites (S) and parsimony informative sites (P), haplotype diversity (Hd), nucleotide diversity (π), and average number of nucleotide differences (k), were estimated using DnaSP. This was done for the whole data set (ingroup), for the main phylogroups, and also for other relevant geographic groups, including island populations and continental samples.

### Population genetic structure

Pairwise genetic differentiation values (F_ST_) between all pairs of phylogroups and other relevant geographic groups, and an Analysis of Molecular Variance (AMOVA) were calculated using Arlequin version 3.11 (Excoffier *et al*., 2005). Ten thousand nonparametric permutations were performed to generate a random distribution to test the significance of the pairwise F_ST_ values and covariance components of the AMOVA, and α = 0.05 was set as the threshold for statistical significance.

A Mantel test was used to evaluate the relationship between matrices of pairwise geographic distances and genetic differentiation values (Slatkin’s linearised pairwise F_ST_ as D = F_ST_/(1-F_ST_); Slatkin, 1995). Despite criticisms, the Mantel test is still a widely used and can be a powerful statistical approach to analyse sequence data to test evolutionary hypotheses (Diniz-Filho *et al*., 2013). Due to the very low (or absence of) genetic variation in the Orkney islands, DNA sequences originating from there were pooled to avoid issues with pairwise F_ST_ calculations.

Geographic barriers were computed using Barrier version 2.2 (Manni *et al*., 2004). This approach implements Monmonier’s maximum difference algorithm to find edges (boundaries) on a Voronoi tessellation associated with the highest rate of change in genetic distances among samples interconnected by a geometric network (i.e. Delaunay triangulation) (Manni *et al*., 2004). A barrier highlights the geographic areas where a genetic discontinuity is found, and where samples on each side of the barrier are genetically more similar than samples taken on different sides of the boundary. Pairwise genetic distances were estimated using continental samples only, limiting the data set in the geometric network calculation to one sample per locality, and computing a maximum of 10 barriers.

### Historical demography

A strict molecular clock was compared to the uncorrelated lognormal relaxed molecular clock (Drummond *et al*., 2006). Coalescent constant population size and Bayesian skyline demographic models (Drummond *et al*., 2005) were compared to identify the best-fitting pattern of changes in the pygmy shrew population. For model selection, path sampling and stepping-stone sampling (Baele *et al*., 2013), based on four independent MCMC chains (1000 steps of 100,000 generations each, following a 10 million generations burn-in period), were used for calculating the log Marginal Likelihoods Estimates (MLEs) for each model. MLEs were used to calculate Bayes Factors (BFs) for each comparison between tested models to determine the best-fitting one (Kass & Raftery, 1995). The best-fitting models were then used to estimate the Time of divergence from the Most Recent Common Ancestor (TMRCA) and Bayesian Skyline Plots (BSP) (see below). The 95% Highest Posterior Density (HPD) was included in the TMRCA and BSP estimations.

TMRCAs for the ingroup (all *S. minutus* samples) and the phylogroups were estimated using BEAST version 2.5.2 (Bouckaert *et al*., 2014). The following prior assumptions were: random starting tree, monophyletic groups (for the ingroup and the Irish phylogroups) (Drummond *et al*., 2006) to calculate the evolutionary rate, and the GTR substitution model with four categories, gamma = 0.9680 and proportion of invariable sites = 0.4680 (from jModelTest using the full data set). The oldest record of *S. minutus* has been found in Podlesice and Mała Cave, Poland dated between 5.3 and 3.6 MYA (Early Pliocene; Mammal Neogene 14) (Rzebik-Kowalska, 1998). Using this fossil information, a calibration point for the ingroup was set at 4.45 MYA (SD = 0.5 MY; 5.27 – 3.63 MYA) with a normal prior distribution. A second calibration was set for the Irish lineage at 0.006 MYA (SD = 0.0005 MY; 0.00682 – 0.00518 MYA) based on the inferred colonisation time of Ireland by *S. minutus* using control region sequences (McDevitt *et al*., 2009). The trace files were analysed in Tracer, the tree information from the four runs were combined and resampled at a lower frequency (for a total of 10,000 trees) using LogCombiner, and the information was summarized using TreeAnnotator selecting Maximum clade credibility tree and median heights. The phylogenetic tree showing the TMRCAs was created using FigTree version 1.4.4 (http://tree.bio.ed.ac.uk/software/figtree/).

Genetic evidence of population expansion for the phylogroups, island populations and continental samples was investigated using the R_2_ test of neutrality (Ramos-Onsins & Rozas, 2002), based on the difference of the number of singleton mutations and the average number of nucleotide differences, and Fu’s Fs (Fu, 1997), a statistic based on the infinite-site model without recombination that shows large negative Fs values when there has been a demographic population expansion. Both population expansion tests were carried out in DnaSP using coalescent simulations for testing significance (10,000 replicates).

Mismatch distributions (i.e. the distribution of the number of differences between pairs of haplotypes) were estimated for the phylogroups (and where N ≥ 10) to compare the demography of the populations with the expectations of a sudden population expansion model (Rogers & Harpending, 1992). For the phylogroups and continental samples that showed a unimodal mismatch distribution and significant population expansion, the time since the population expansion (t) was calculated as t = τ/2u, where τ (tau) is the mode for the unimodal mismatch distribution, and u is the cumulative (across the sequence) probability of substitution (Schenekar & Weiss, 2011). The calculations were done using the MS Excel Mismatch Calculator (Schenekar & Weiss, 2011) with sequence length = 1110 bp, generation time = 1 year (Hutterer *et al*., 2016), percent divergence/MY = 0.551 (based on the average substitution rate across all sites clock rate results from BEAST) and cumulative substitutions/generation = 0.00062.

BSPs were calculated using BEAST based on the posterior distribution of effective population size through time from a sample of gene sequences. This was done for the phylogroups showing a unimodal mismatch distribution and significant signatures of recent population expansion (where N ≥ 10). The analysis was run for 100 million generations, sampled every 1000, using the best-fitting model.

## RESULTS

### Phylogenetic analysis

For the complete *S. minutus* data set (N = 671) (Fig. 1B), there were 424 haplotypes with 390 polymorphic sites of which 277 were parsimony informative (Table 1). We report 160 newly sequenced specimens of *S. minutus* from the Iberian (4) and Balkan (19) peninsulas and from Central and Northern Europe (137) from which 127 were new haplotypes. Also, there were three new sequences and haplotypes of *S. volnuchini*, from which two were from Turkey and one from the Crimean Peninsula.

**Table 1.** DNA sequence polymorphism in *Sorex minutus* phylogroups and other geographic groups

| Phylogroups | N | S | P | H | Hd | Hd (SD) | $\pi$ | $\pi$ (SD) | k |
| --- | --- | --- | --- | --- | --- | --- | --- | --- | --- |
| Ingroup | 671 | 390 | 277 | 424 | 0.9899 | 0.0015 | 0.0143 | 0.0000 | 15.8670 |
| Italian | 26 | 51 | 19 | 18 | 0.9600 | 0.0230 | 0.0061 | 0.0004 | 6.7720 |
| South Italian | 4 | 16 | 0 | 4 | 1.0000 | 0.1770 | 0.0072 | 0.0020 | 8.0000 |
| Balkan | 22 | 55 | 28 | 17 | 0.9610 | 0.0290 | 0.0097 | 0.0009 | 10.7970 |
| Iberian | 6 | 15 | 6 | 5 | 0.9330 | 0.1220 | 0.0058 | 0.0013 | 6.4000 |
| Western | 283 | 147 | 83 | 102 | 0.9458 | 0.0067 | 0.0049 | 0.0002 | 5.4400 |
| Irish | 94 | 53 | 21 | 42 | 0.8920 | 0.0270 | 0.0020 | 0.0002 | 2.2180 |
| Northern | 330 | 311 | 197 | 278 | 0.9984 | 0.0005 | 0.0065 | 0.0002 | 7.1840 |
| Continental groups |  |  |  |  |  |  |  |  |  |
| Western (Continental) | 15 | 28 | 11 | 13 | 0.9810 | 0.0310 | 0.0050 | 0.0006 | 5.5430 |
| Northern (Continental) | 226 | 241 | 142 | 188 | 0.9978 | 0.0007 | 0.0062 | 0.0002 | 6.9300 |
| Other island groups |  |  |  |  |  |  |  |  |  |
| Orkney Islands (All) | 119 | 17 | 13 | 11 | 0.7720 | 0.0210 | 0.0027 | 0.0001 | 3.0140 |
| Orkney Mainland | 44 | 9 | 7 | 8 | 0.7550 | 0.0550 | 0.0013 | 0.0002 | 1.4790 |
| Orkney South Ronaldsay | 40 | 1 | 1 | 2 | 0.1420 | 0.0710 | 0.0001 | 0.0001 | 0.1420 |
| Orkney Westray | 33 | 0 | 0 | 1 | 0.0000 | 0.0000 | 0.0000 | 0.0000 | 0.0000 |
| Orkney Hoy | 2 | 2 | 0 | 2 | 1.0000 | 0.5000 | 0.0018 | 0.0009 | 2.0000 |
| Belle Île | 5 | 9 | 3 | 5 | 1.0000 | 0.1260 | 0.0038 | 0.0010 | 4.2000 |
| British | 91 | 146 | 61 | 80 | 0.9960 | 0.0030 | 0.0055 | 0.0003 | 6.1210 |

N = Sample size; S = Number of polymorphic (segregating) sites; P = Parsimony informative sites; H = Number of haplotypes; Hd = Haplotype diversity; SD = Standard Deviation; $\pi$ = Nucleotide diversity; k = Average number of nucleotide differences.

The Bayesian phylogenetic analysis showed *S. minutus* as a monophyletic group and revealed six distinct lineages corresponding to their geographical origin (i.e. phylogroups) supported by high posterior probabilities (Fig. 2A). Samples from the Mediterranean peninsulas clustered in three distinct phylogroups, namely the Iberian, Italian and Balkan phylogroups. The Iberian group was represented with few DNA sequences (N = 6). It was geographically restricted to the Iberian Peninsula and included samples from Rascafría, Central Spain (Sierra de Guadarrama) and Picos de Europa, Northern Spain. The Italian phylogroup (N = 26) was mostly restricted to the north-central regions of the Italian peninsula; it included samples from the Apennines and the Alps in Italy, but also from Switzerland, Slovenia, Southern and Eastern France near the border with Italy, Czech Republic and Germany. The Balkan phylogroup (N = 22) included samples mostly from the Balkan Peninsula and a few from further north in Central Europe. This phylogroup showed a weak north/south subdivision, with one clade containing samples from Switzerland, Austria, Slovakia, Czech Republic, Hungary and Montenegro, another clade containing samples from Serbia, Bosnia and Herzegovina and North Macedonia, plus other ungrouped basal samples from Montenegro, North Macedonia, Serbia and Turkey (East Thrace, Southeast Europe).

**Figure 2.**
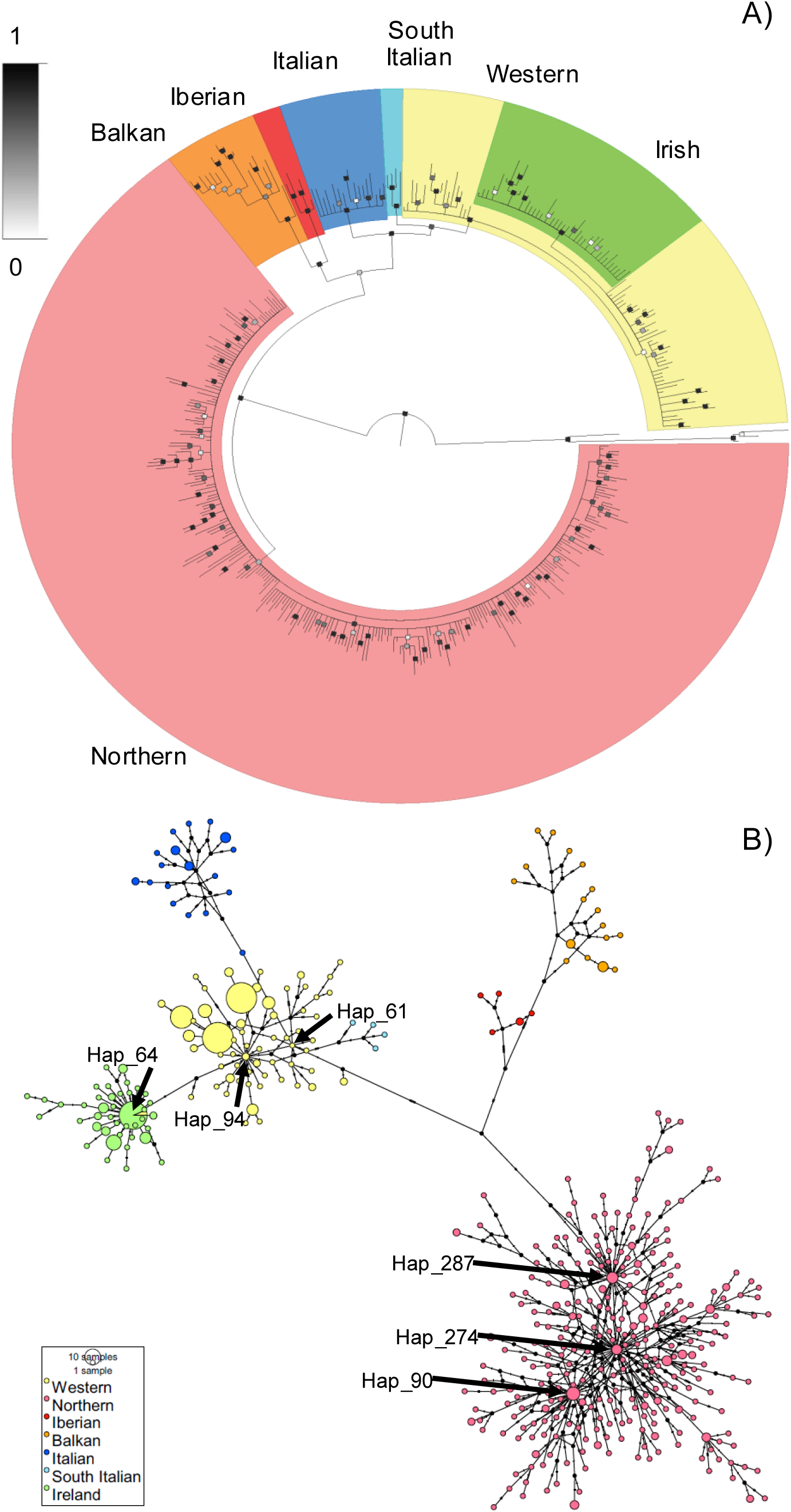
Phylogenetic reconstructions of the Eurasian pygmy shrew *Sorex minutus* using cyt b sequences. A) Bayesian phylogenetic tree (with posterior probabilities on branches) showing the phylogroups. B) Haplotype phylogenetic network with haplotypes represented as nodes and their evolutionary relationships represented by edges; relevant haplotypes named at the centre of star-like patterns.

There was also a well-supported and geographically widespread Western phylogroup (N = 283), which included samples from northern Spain (Cantabrian Mountain Range), Southern France and Andorra (i.e. the Pyrenees), western and central France (including Belle-Île), Ireland, the Orkney Islands, and western mainland Britain and offshore islands on the western coast of mainland Britain. Samples from Ireland formed an internal monophyletic lineage (i.e. the Irish phylogroup, N = 94) within the Western phylogroup. Notably, two samples from Navarra in northern Spain (ESNa0861 and ESNa1131; Accession Number JF510331) shared haplotypes with samples from Ireland. A monophyletic South Italian phylogroup (N = 4) was most closely related to the Western phylogroup than to the Italian phylogroup, and was geographically restricted to La Sila, Calabria in Southern Italy.

Samples from northern and central Europe and Siberia, namely the Northern phylogroup (N = 330), formed the most geographically widespread lineage and included samples ranging from Central France and Britain (excluding those within the Western phylogroup), across Central and Northern Europe to Lake Baikal in Siberia, but did not include samples from Southern Europe. Samples from mainland Britain belonging to the Northern phylogroup did not form an internal monophyletic cluster.

The phylogenetic network had a complex structure (Fig. 2B), but the haplotypes clustered into the same phylogroups detected with Bayesian phylogenetics and were distantly related from each other (> 10 mutational steps). The Western phylogroup had a star-like pattern and showed three most internal haplotypes; notably, one internal haplotype (Hap_64) included samples from Northern Spain and Ireland, and many other Irish haplotypes derived from it. The Northern phylogroup showed a star-like pattern with many reticulations and three most internal haplotypes separated from each other by few mutational steps. There was an apparent geographical subdivision within the Northern phylogroup, where samples from Siberia, Eastern and Northern Europe were derived or most closely connected to samples from Central Ukraine (Hap_287), samples from Central Europe were derived or most closely connected to samples from The Netherlands (Hap_274), and all samples from Britain were derived or most closely connected to other samples from The Netherlands than to the other central haplotypes (Hap_90); however, the highly reticulated pattern of the inner haplotypes of the Northern phylogroup indicated that this geographical subdivision was weak.

Sequence polymorphism indices and diversity values for the phylogroups and other geographic groups are shown in Table 1. For the phylogroups, the haplotype diversity values were high (>90%), and the nucleotide diversity values were either half or almost half as much as the ingroup. Notably, the Northern phylogroup had the highest haplotype diversity values, followed by the Balkan phylogroup; however, the Balkan phylogroup had the highest nucleotide diversity values. The Irish phylogroup, which clustered within the Western phylogroup, showed slightly lower haplotype diversity than any other phylogroups.

The continental groups (Northern continental and Western continental) showed equivalent DNA polymorphism values as the main phylogroups, but the island groups showed different levels of DNA polymorphism (Table 1). There was low DNA polymorphism in islands of the Orkney Archipelago, with only 11 haplotypes in all Orkney Islands combined (N = 119), but all haplotypes were unique to these islands. There were eight haplotypes in Orkney Mainland (N = 44), from which seven were unique to this island (the largest island of the archipelago), there were two unique haplotypes in Orkney South Ronaldsay (N = 40), and there was only one haplotype in Orkney Westray (N = 33) also present in Orkney Hoy (N = 2) and Orkney Mainland. There were five haplotypes in Belle-Île (N = 5), and only one was present in the continent also belonging to the Western phylogroup. The British group (N = 91) showed high haplotype diversity but moderate nucleotide diversity values and had 80 haplotypes from which 77 were unique haplotypes not found elsewhere.

### Population genetic structure

The highest pairwise differentiation values were found between some southern phylogroups and island groups, while the lowest values were between phylogroups and islands groups that clustered within them (Supplementary information Table S2). There was higher percentage of variation among (73.5 %) than within (26.5 %) groups, and there was a significant population differentiation (F_ST_ = 0.7349, P < 0.0001). The Mantel test showed a nonsignificant relationship between pairwise geographic and genetic distances based on Slatkin’s linearised F_ST_ (R_2_ = 0.0095, P = 0.2935) (Supplementary information Fig. S1).

The barriers identified using the computational geometry approach reflected the genetic differentiation between *S. minutus* and *S. volnuchini*, and among the phylogroups within *S. minutus* (Fig. 1C). The first barrier separated *S. minutus* from *S. volunichini*. The nine following barriers coincided with the location of mountain ranges, including a barrier located in the north of the Balkan Peninsula, in the Alps and in the Pyrenees, which reflected the genetic subdivisions and lineages in *S. minutus*.

### Historical demography

Comparison of BFs for each model indicated the Bayesian skyline demographic model as the best-fitting one (BF = 391), and the strict molecular clock was better than the uncorrelated lognormal relaxed molecular clock (BF = 23). The MLEs for the constant population size and Bayesian skyline demographic models using the strict molecular clock were -10960 and -10569, while using the uncorrelated lognormal relaxed molecular clock were -10907 and -10592, respectively. Therefore, the strict clock and Bayesian skyline demographic model were selected as the best-fitting according to BFs. The effective sample size (ESS) for all values was higher than 200.

All branches of the Bayesian genealogy (Fig. 3, Table 2) were well-supported (posterior probabilities PP ≥ 0.97), except for the clade containing all phylogroups excluding Iberian (PP = 0.05). The ingroup split approximately KYA 83.4, with lower and upper 95% highest posterior density HPD limits of approximately between 59.7 and 110.2 KYA. The Iberian phylogroup split approximately 31.8 KYA (95% HPD: 22 – 43.1 KYA, respectively. The Balkan phylogroup had a TMRCA of approximately 29.6 KYA (95% HPD: 21.8 – 40.5 KYA). The Northern and Western phylogroups split approximately 24.1 KYA (95% HPD: 16.4 – 33.1 KYA), and the Irish phylogroup arose approximately 5.9 KYA (95% HPD: 4.9 – 6.9 KYA). The Italian phylogroup had a TMRCA of approximately 15.3 KYA (95% HPD: 10.7 – 21.5 KYA), while the South Italian phylogroup of approximately 12.8 KYA (95% HPD: 8.5 – 17.8 KYA).

**Figure 3.**
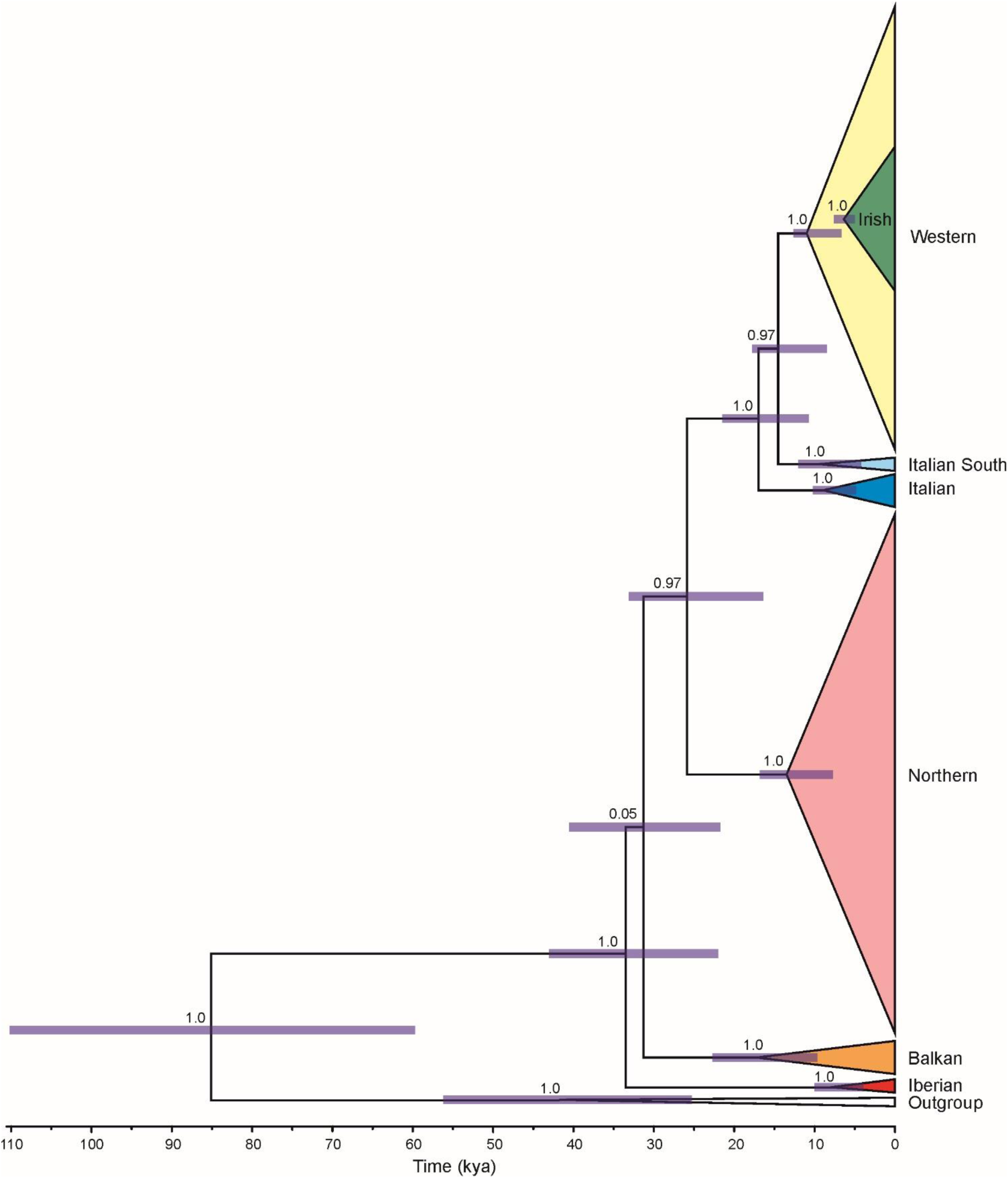
Time of divergence from the Most Recent Common Ancestor (TMRCA) for the main phylogroups. Numbers on nodes represent posterior probabilities, and horizontal bars represent the 95% Highest Posterior Density (HPD). Dates in Thousand Years Ago (KYA).

**Table 2.** Population expansion tests for *Sorex minutus* phylogroups and other geographic groups

| Phylogroups | R<sub>2</sub> | P-value | F<sub>s</sub> | P-value | τ | t<br>(in years) | TMRCA<br>(in KYA) | 95% HPD<br>(in KYA) |
| --- | --- | --- | --- | --- | --- | --- | --- | --- |
| Ingroup | 0.0198 | **0.0004** | -741.2620 | *** | 7.8590 | 6425 | 83.4 | 59.7-110.2 |
| Italian | 0.0521 | **0.0000** | -5.8766 | **0.0152** | 6.7720 | 5536 | 15.3 | 10.7-21.5 |
| South Italian | 0.1822 | 0.1658 | 0.0687 | 0.2975 | 5.6340 | - | 12.8 | 8.5-17.8 |
| Balkan | 0.0830 | 0.0542 | -3.6701 | 0.0768 | 7.1500 | - | 29.6 | 21.8-40.5 |
| Iberian | 0.1462 | 0.0888 | 0.0731 | 0.4290 | 4.0100 | - | 31.8 | 22.0-43.1 |
| Western | 0.0175 | **0.0004** | -114.6990 | *** | 3.6660 | 2997 | 24.1 | 16.4-33.1 |
| Irish | 0.0187 | **0.0000** | -52.5664 | *** | 1.3040 | 1066 | 5.9 | 4.9-6.9 |
| Northern | 0.0105 | **0.0000** | -663.4730 | *** | 6.5390 | 5346 | 24.1 | 16.4-33.1 |
| Continental groups |  |  |  |  |  |  |  |  |
| Western (Continental) | 0.0793 | **0.0045** | -6.0342 | **0.0035** | 5.5430 | 4532 | - | - |
| Northern (Continental) | 0.0128 | **0.0000** | -386.4520 | *** | 5.8010 | 4742 | - | - |
| Other island groups |  |  |  |  |  |  |  |  |
| Orkney Islands (All) | 0.0880 | 0.5209 | 0.6044 | 0.6437 | 1.1740 | - | - | - |
| Orkney Mainland | 0.0839 | 0.2301 | -1.6879 | 0.1892 | 1.4790 | - | - | - |
| Orkney Hoy | 0.5000 | 1.0000 | NC | NC | 2.0000 | - | - | - |
| Orkney South Ronaldsay | 0.0712 | 0.1770 | -0.2182 | 0.4420 | 0.1420 | - | - | - |
| Orkney Westray | NC | NC | NC | NC | NC | - | - | - |
| Belle Île | 0.1915 | 0.2467 | -1.6330 | 0.0732 | 3.5500 | - | - | - |
| British | 0.0161 | **0.0000** | -122.8550 | *** | 6.1210 | 5004 | - | - |

R<sub>2</sub> = Ramos-Onsins and Rozas (2002) test of neutrality; P-value = P-values of expansion tests expected under neutrality (\*\*\* = P < 0.001); F<sub>s</sub> = Demographic population expansion test (Fu 1997); τ = (2ut) The mode of a mismatch distribution; t = Time of population expansion (for phylogroups with bi- or unimodal mismatch distributions); TMRCA = Time of divergence from the Most Recent Common Ancestor; 95% HPD = 95% Highest Posterior Density; KYA = Thousand Years Ago; NC = Not computable (not enough variation or samples)

The population expansion tests (R_2_ and Fu’s Fs) showed significant departures from neutrality for the ingroup and several other phylogroups, except for the Balkan, Iberian and South Italian (Table 2). The population expansions were not an effect of the island samples belonging to these phylogroups, and continental samples analysed separately also demonstrated a similar pattern (Table 2). For the island groups, only the Irish and British groups showed signatures of recent population expansions (Table 2).

The mismatch distributions varied significantly among the phylogroups (Fig. 4A; Supplementary information Fig. S2). The ingroup showed a bimodal mismatch distribution, which reflected the pairwise comparisons within and among phylogroups in *S. minutus*. The Northern (and Northern continental), Italian, Western (and Western continental) and Irish phylogroups all had distinctly unimodal distributions with an almost perfect fit between observed and expected pairwise differences of a sudden population expansion model. All population expansions for the phylogroups were dated to the Holocene; the Italian and Northern phylogroups had the oldest times of expansion (>8.0 KYA), while the Irish showed a relatively recent population expansion dated to 1.6 KYA.

**Figure 4.**
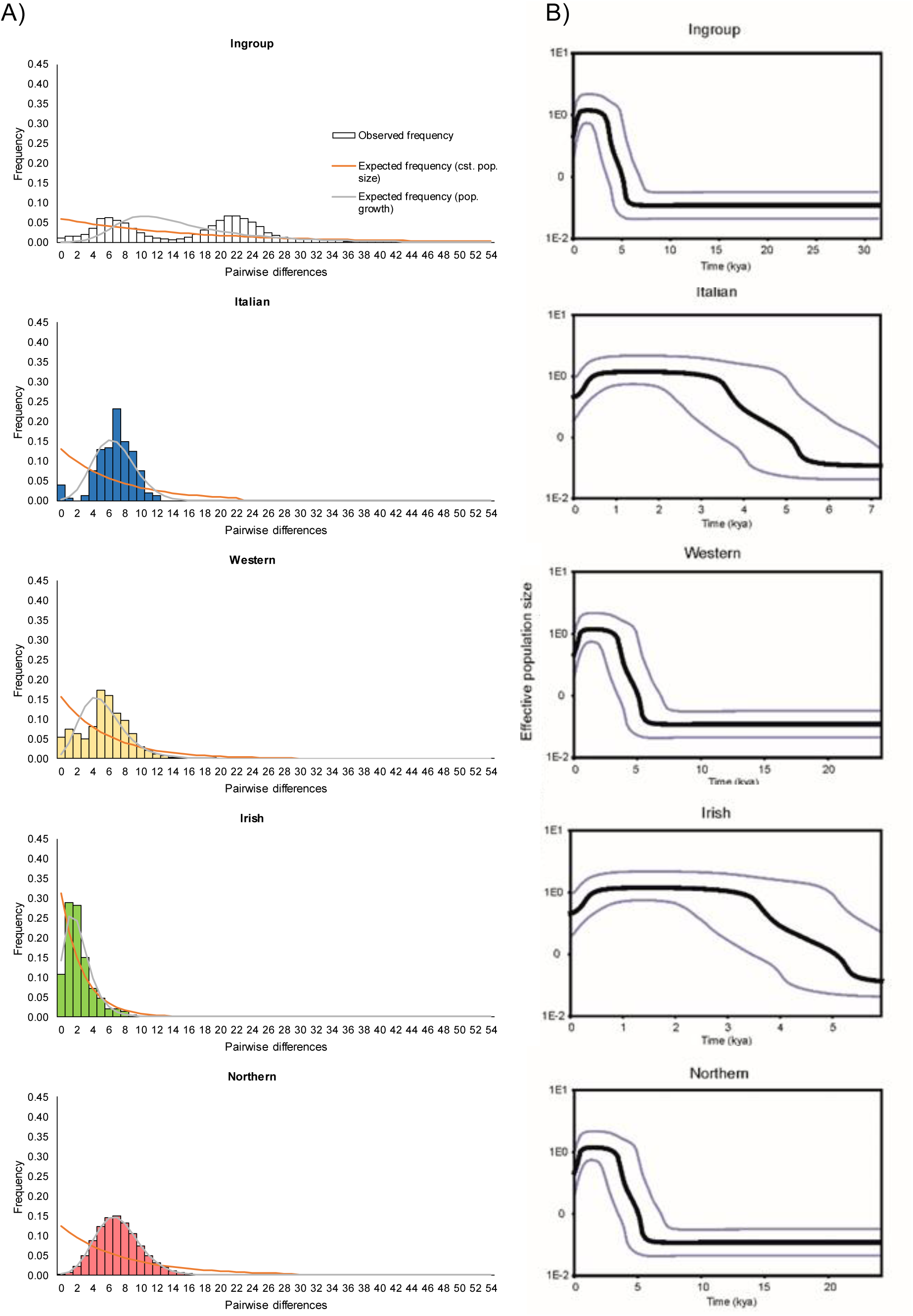
Historical demography of the Eurasian pygmy shrew *Sorex minutus*. A) Mismatch distributions of groups with significant signatures of population expansion. B) Bayesian Skyline Plots (BSP) of phylogroups with significant signatures of population expansion. The solid lines in BSP are median estimates and the shaded areas represent 95% Highest Probability Densities (confidence intervals).

The BSP obtained for three phylogroups (Northern, Western and Irish) suggested that demographic expansions of these populations started approximately 5.0 KYA (Fig. 4B). BSP calculation for the Italian phylogroup indicated an even earlier demographic expansion (approximately 5.5 KYA) (Fig. 4B).

## DISCUSSION

Quaternary refugia represent the geographical regions that species inhabit during periods of glacial and interglacial cycles when there is the maximum contraction in geographical range (Stewart *et al*., 2009). There is support for both southern (Taberlet *et al*., 1998; Hewitt, 2000) and northern glacial European refugia (Bilton *et al*., 1998; Stewart & Lister, 2001; Kotlík *et al*., 2006; Provan & Bennett 2008; Fløjgaard *et al*., 2009; Vega *et al*., 2010a, b). Rather than polarising the biogeographic patterns into southern and northern refugia (Tzedakis *et al*., 2013), the paradigms of postglacial colonisation in Europe (Hewitt, 2000) can be improved with the acceptance of southern hotspots of diversification without northward colonisation (Bilton *et al*., 1998) and the concept of refugia-within-refugia (Gómez & Lunt, 2007), as well as with the findings of cryptic northern glacial refugia (Stewart & Lister 2001; Provan & Bennett, 2008; Stewart *et al*., 2009), to reflect the evolutionary processes across varied topographical areas that have shaped genetic diversity. The statistical phylogeographic results obtained here, using published and newly described samples and haplotypes, notably expand previous findings on *S. minutus*, giving a more precise population genetic structure and demographic history. Thus, our findings on *S. minutus* contribute to the understanding of the phylogeographic patterns and processes during the Quaternary glaciations that shaped the European biota, and contribute to the emerging pattern of complex biogeographical histories in Europe (Pedreschi *et al*., 2019).

### Sorex minutus *phylogeography*

The significant genetic structure among phylogroups defined in this study illustrate the complex history of European colonisation, isolation and diversification of *S. minutus* during the Pleistocene and Holocene, and is not a simple case of isolation by distance and colonisation of Northern and Central Europe from expanding populations from the south. While the southern phylogroups, including the Iberian, Balkan, Italian and South Italian, were mostly restricted to the Southern European peninsulas (consistent with the traditional southern glacial refugia), the Northern and Western phylogroups were widespread geographically and were found north of the Mediterranean peninsulas, consistent with previous studies with fewer samples (Bilton *et al*., 1998; Mascheretti *et al*., 2003; Vega *et al*. 2010a, b) and with different molecular markers (McDevitt *et al*., 2010).

The hypothesis of northern refugia is further supported by palaeontological and palynological evidence for other temperate and boreal species (Willis *et al*., 2000; Willis & van Andel, 2004; Magri *et al*., 2006; Sommer & Nadachowski, 2006), as well as many phylogeographic studies in small mammals, including the field vole *M. agrestis* (Jaarola & Searle, 2002), bank vole *M. glareolus* (Deffontaine *et al*., 2005; Kotlík *et al*., 2006; Wójcik *et al*., 2010), root vole *M. oeconomus* (Brunhoff *et al*., 2003), common vole *M. arvalis* (Heckel *et al*., 2005; Stojak *et al*., 2016), common shrew *S. araneus* (Bilton *et al*., 1998; Yannic *et al*., 2008) and weasels *Mustela nivalis* (McDevitt *et al*., 2012). For several small mammals, including *S. minutus*, suitable climatic conditions at the LGM could have been widespread across Central and Eastern Europe (Fløjgaard *et al*., 2009; Vega *et al*., 2010b; McDevitt *et al*. 2012; Stojak *et al*., 2019).

Until recently, it was unclear which species of *Sorex* inhabit Crimea. According to Zagorodniuk (1996) it could be *S. (minutus) dahli* [mentioned in Hutterer (2005) as a synonym of *Sorex volnuchini* (*dahli*)], and Zaitsev *et al*. (2014) and Hutterer *et al*. (2016) showed *S. minutus* in mainland Ukraine and in Crimea. Hutterer (2005) mentioned that *S. volnuchini* might be present in Crimea, but in Hutterer *et al*. (2016) *S. volnuchini* is only shown in southern Russia and Caucasus States, Turkey and northern Iran. Our research demonstrated that *S. volnuchini* may be present in the southern region of Crimea (based on one cyt b sequence), while *S. minutus* is present in the mainland, but further sampling in this region is needed.

The finding that in both the Iberian and Italian peninsulas there are two genetic lineages of *S. minutus* (four in total) suggests that the refugial areas may have had subdivisions at the LGM. In the Iberian Peninsula, the topography of the region with east- west mountain ranges and other high ground (over 1000 m a.s.l.), large rivers (which could act as barriers to dispersal), and the distinct seasonal precipitation and vegetation types (O’Regan, 2008), must have played an important role in the genetic differentiation of populations and could explain the presence of two phylogroups (i.e. the Iberian and Western phylogroups). McDevitt *et al*. (2010) proposed that the Western phylogroup could have originated in the Dordogne region based on a limited number of samples from France but the presence of this phylogroup in northern Iberia could mean that an Iberian origin is possible instead. A similar process could explain the presence of the two phylogroups in the Italian peninsula (i.e. Italian and South Italian). The genetic differentiation of the South Italian phylogroup, further supported by morphological data (Vega *et al*., 2010a), could be due to the unique geography of Southern Italy consisting of mountain massifs of Pollino, La Sila and Aspromonte separated by lowland areas, which from the Pliocene to the end of the Middle Pleistocene, at times of high sea level, were islands in a chain (Malatesta, 1985; Caloi *et al*., 1989; Bonardi *et al*., 2001; Bonfiglio *et al*., 2002). The patterns of differentiation within refugial areas were concordant with the ‘refugia-within-refugia’ concept widely recognized for the Iberian Peninsula (Gómez & Lunt, 2007; Abellán & Svenning, 2019) and similar to microrefugia in the Balkans (Kryštufek *et al*., 2007). For the Italian peninsula, a comparable ‘refugia-within-refugia’ pattern was found in several species (Amori *et al*., 2008; Canestrelli *et al*., 2008; Castiglia *et al*., 2008; Vega *et al*., 2010a; Senczuk *et al*., 2017; Bisconti *et al*., 2018).

The genetic similarity between the Western and South Italian phylogroups indicates a common history and it can be hypothesised that their common ancestor was more widespread throughout the Italian peninsula, probably displaced later by the Italian lineage in the Apennines and Western Alps. A similar scenario has been proposed for the water shrew *Neomys fodiens* (Castiglia *et al*., 2007), Alpine salamander *Salamandra salamandra* (Steinfartz *et al*., 2000), black pine *Pinus nigra* (Afzal-Rafii & Dodd, 2007) and green lizard *Lacerta bilineata bilineata* (Böhme *et al*., 2007), which showed closely related South Italian and Western phylogroups most closely related to each other than to a North-Central Italian lineage.

The phylogeographic patterns found here were further supported by the determination of barriers that coincided with mountain ranges located on the north of the Iberian, Italian and Balkan peninsulas. Contact zones among phylogroups (i.e. localities where at least two cyt b phylogroups were present) were detected at the northern extremes of the southern peninsulas. During the LGM, glaciers covered most of the Alpine (Buoncristiani & Campy, 2004) and Pyrenean mountain ranges (Calvet, 2004), while glaciers in the Carpathians (Reuther *et al*., 2007) and in the Balkan Peninsula (Hughes *et al*., 2006) were found > 1,000 m a.s.l. When climate ameliorated and suitable habitat became available, pygmy shrew populations belonging to different phylogroups on different sides of the mountain ranges could have expanded and colonised previously glaciated areas thus forming the observed contact zones. Moreover, the widespread distribution and absence of phylogeographic structure of the Northern phylogroup could be explained by the apparent absence of major geographical barriers across Central and Northern Europe, and recolonization from northern refugia. Similarly, pygmy shrews belonging to the Western and Northern phylogroups could have quickly colonised mainland Britain across a land connection to continental Europe (i.e. Doggerland; Gaffney *et al*., 2007), resulting in the genetic similarities observed between the British Isles and continental Europe.

### Sorex minutus *demography*

The oldest fossils assigned to *S. minutus* were found in Podlesice and Mała Cave, Poland dated to the Early Pliocene between 4 and 5.3 MYA (Rzebik-Kowalska, 1998). An early widespread colonisation of Europe by *S. minutus* might have been possible because it was probably one of the first species of the genus *Sorex* in the continent (Rzebik-Kowalska, 1998, 2008). The Bayesian analysis revealed, however, recent timing of diversification events, with TMRCAs for the ingroup and the phylogroups in continental Europe between the Upper Pleistocene and Lower Holocene, and in the Middle Holocene for the Irish phylogroup. Similar colonisation scenarios and divergence before the LGM from Eastern to Western Europe have been proposed for other species, including the common vole *Microtus arvalis* (Heckel *et al*., 2005; Stojak *et al*., 2016), the bank vole *Clethrionomys glareolus* (Deffontaine *et al*., 2005; Kotlík *et al*., 2006; Wójcik *et al*., 2010), and the root vole *M. oeconomus* (Brunhoff *et al*., 2003).

The population expansion signatures for the Northern and Western phylogroups, star-like patterns in phylogenetic networks and population expansion times support recent and quick colonisation events of central and northern Europe, and appear to reflect responses to postglacial climate warming. The Western lineage was restricted to Central, Western and South-Eastern France and North-Western Spain in continental Europe, but it was the only lineage found in Ireland and several islands off the west and north coasts of Britain. The region of the Dordogne in South-Western France was situated outside the LGM permafrost area and has temperate mammal fossil records dated to the end of the LGM; therefore, it has been suggested as another likely northern refugium (Sommer & Nadachowski, 2006; McDevitt *et al*., 2010) where the Western lineage could have persisted and recolonised Western and Central France after the LGM. But as stated above, an Iberian origin for this phylogroup is also possible. However, SDM studies showed that suitable climatic conditions during the LGM for *S. minutus* and other temperate small mammal species could have been more continuous and present further north (Fløjgaard *et al*., 2009; Vega *et al*., 2010b), which could explain its widespread distribution in Western Europe and its presence in Britain. According to BSP results, it is plausible that Northern and Western phylogroups spread across Europe after the Younger Dryas (11.7 to 12.9 KYA). The British (island) group, belonging to the Northern phylogroup, showed a significant signature of population expansion. This expansion could have selectively displaced pygmy shrew populations of the Western lineage, which still remain in uplands and islands in the periphery to the north, west and south of Britain forming a ‘Celtic fringe’ (Searle *et al*., 2009).

The widespread Italian lineage may be presumed to derive from a glacial refugium located somewhere within the vicinity of the Apennine mountain chain. A significant population expansion signature demonstrates that the Italian phylogroup went through a recent expansion phase, calculated in BSP for about 5.5 KYA. Contrastingly, the lack of a population expansion signature, the high nucleotide and haplotype diversities, and the highly divergent sequences showing a weak north/south subdivision of the Balkan phylogroup warrants further attention. The Balkans is a European hotspot for biodiversity given its environmental stability, topographic and climatic diversity and occasional connectedness with Asia Minor (Kryštufek & Reed, 2004; Kryštufek *et al*., 2007, 2009; Bužan *et al*., 2010), and it could be expected that some of these factors shaped the genetic diversity of the Balkan lineage there. Similarly, the lack of significant population expansion values for the Iberian lineage may relate to historical stable population sizes; however, the sample size was low and this result should be taken with caution.

### Further considerations and implications

The comparison of the results obtained here with those elsewhere shows an emerging pattern of glacial refugia in Mediterranean peninsulas and further north in Central Europe for several species.

Although *S. minutus* is considered as a least concern species by the IUCN (Hutterer *et al*., 2016), the distinct phylogroups deserve more attention than this implies. Genetic diversity is considered an important aspect of global biodiversity (McNeely *et al*., 1990), and local and/or country-based conservation efforts are highly valued (for example, in Britain and Ireland the pygmy shrew is protected by law). The refugial areas in Southern Europe are often found in mountain ranges at the low-latitude margins of the present-day distribution ranges of species and are most likely to contain rear-edge populations where selection for local adaptations could have resulted in the evolution of distinct ecotypes (Cook, 1961; Hampe & Petit, 2005). Rear-edge populations, including the southern lineages of *S. minutus*, deserve further investigation and should be regarded for conservation because they are important to determine the responses of species to modern climate change (Petit *et al*., 2003; Hampe & Petit, 2005).

In conclusion, the Eurasian pygmy shrew *Sorex minutus* is a good model for understanding biological diversity, colonisation patterns and the effects of past climate change on biological diversity. There is a mosaic of genetic lineages across continental Europe, characterised by different demographic histories and natural colonisation patterns, while island populations are characterised by recent natural and human-mediated colonisations. This study has notably expanded previous findings on *S. minutus*, with a more precise statistical phylogeographic analysis of the genetic variability and structure, colonisation routes, geographical barriers and historical demography across Europe. Specifically, we provided new data from the Iberian and Balkan peninsulas, and from Central and Eastern Europe (Poland, Ukraine and Russia), important for understanding postglacial events. *Sorex minutus* is not an easy species to obtain in large numbers, and the sampling described here represents a very substantial effort. However, it is a species that is unusually widespread and genetically subdivided and therefore can inform better than almost any other about the relative importance of southern and northern glacial refugia.

## ACKNOWLEDGEMENTS

Specimens and species records of *Sorex minutus* were made available by several museums and we acknowledge the help of the curators from the following institutions: Département d’écologie et évolution (Université de Lausanne, Switzerland), Univerza na Primorskem and Research Centre of Koper (Slovenia), Natuurhistorisch Museum (Rotterdam, Netherlands), Dipartimento di Ecologia (Università della Calabria, Italy), Museo di Anatomia Comparata and Museo di Zoologia “La Sapienza” (Università di Roma, Italy), Natuurmuseum Brabant (Tilburg, Netherlands) and Mammal Research Institute PAS (Białowieża, Poland). We are very grateful for the tissue samples provided by Glenn Yannic, Jacques Hausser, Jan Zima, Fríða Jóhannesdóttir, Holger Bruns, Peter Borkenhagen, Natália Martínková, Elena Gladilina, Pavel Goldin and Petr Kotlík.

## SUPPORTING INFORMATION

**Table S1.** *Sorex minutus* dataset and sample information

**Table S2.** Pairwise geographic distances (in Km, below diagonal) and genetic differentiation (Slatkin’s F_ST_, above diagonal) among *Sorex minutus* phylogroups and other geographic groups

|  | Italian | South Italian | Balkan | Iberian | Belle Île | Britain | Northern (Continental) | Western (Continental) | Orkney Islands | Irish |
| --- | --- | --- | --- | --- | --- | --- | --- | --- | --- | --- |
| Italian | - | 1.6558 | 3.0387 | 3.6919 | 1.5940 | 2.8852 | 2.3798 | 1.4534 | 2.8673 | 4.0090 |
| South Italian | 773.14 | - | 2.5113 | 3.3869 | 1.3569 | 2.8562 | 2.3204 | 1.1713 | 2.7079 | 4.6820 |
| Balkan | 547.27 | 628.96 | - | 1.9234 | 2.5617 | 3.1494 | 2.8093 | 2.7975 | 6.2456 | 7.0533 |
| Iberian | 1349.26 | 1768.56 | 1880.98 | - | 4.2650 | 3.1850 | 2.6191 | 3.8498 | 7.3804 | 10.8797 |
| Belle Ile | 1162.82 | 1815.47 | 1701.44 | 640.58 | - | 2.7595 | 2.1790 | 0.3345 | 1.2148 | 2.4003 |
| Britain | 1347.34 | 2108.66 | 1795.12 | 1286.79 | 647.11 | - | 0.1449 | 2.6083 | 3.9225 | 4.7265 |
| Northern (Continental) | 1022.36 | 1488.84 | 863.37 | 2218.78 | 1788.28 | 1554.91 | - | 2.0767 | 2.7910 | 3.2035 |
| Western (Continental) | 903.90 | 1444.78 | 1448.34 | 476.18 | 434.81 | 1006.85 | 1742.75 | - | 0.5436 | 1.1193 |
| Orkney Islands | 3476.34 | 3693.42 | 3127.67 | 4679.14 | 4175.54 | 3739.02 | 2477.61 | 4206.50 | - | 1.4635 |
| Irish | 1652.35 | 2396.73 | 2124.06 | 1311.98 | 726.86 | 346.14 | 1897.29 | 1151.96 | 4016.56 | - |

**Figure S1.**
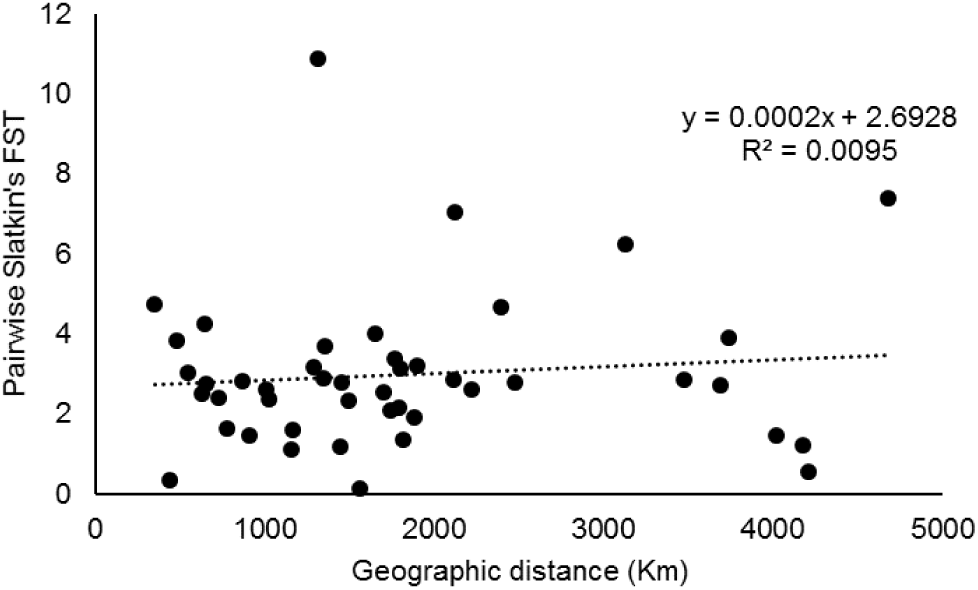
Correlogram of pairwise geographic and genetic distances among *Sorex minutus* phylogroups and other geographic groups.

**Figure S2.**
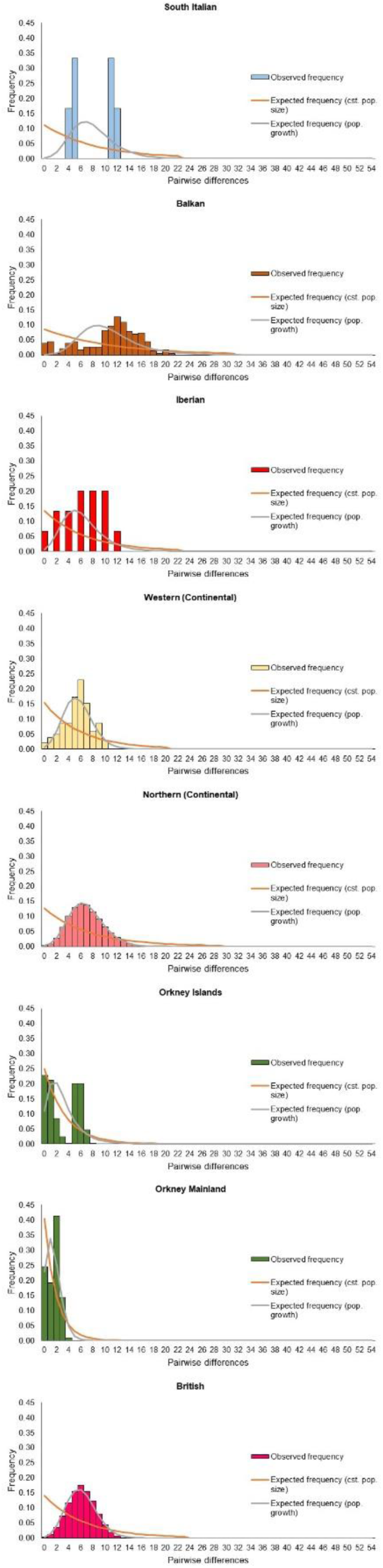
Mismatch distributions of *Sorex minutus* phylogroups and other geographic groups.

